# Universal rules of life: Metabolic rates, biological times and the equal fitness paradigm

**DOI:** 10.1101/2020.07.06.190108

**Authors:** Joseph Robert Burger, Chen Hou, Charles A.S Hall, James H. Brown

## Abstract

Here we review and extend the equal fitness paradigm (EFP) as an important step in developing and testing a synthetic theory of ecology and evolution based on energy and metabolism. The EFP states that all organisms are equally fit at steady state, because they allocate the same quantity of energy, ~22.4 kJ/g/generation to production of offspring. On the one hand, the EFP may seem tautological, because equal fitness is necessary for the origin and persistence of biodiversity. On the other hand, the EFP reflects universal laws of life: how biological metabolism – the uptake, transformation and allocation of energy – links ecological and evolutionary patterns and processes across levels of organization from: i) structure and function of individual organisms, ii) life history and dynamics of populations, iii) interactions and coevolution of species in ecosystems. The physics and biology of metabolism have facilitated the evolution of millions of species with idiosyncratic anatomy, physiology, behavior and ecology but also with many shared traits and tradeoffs that reflect the single origin and universal rules of life.

## Introduction

Life presents a fascinating duality. On the one hand, living things are amazingly diverse. Each of the millions of animal, plant and microbial species is unique. Each has distinctive features of anatomy, physiology, morphology, behavior and ecology that reflect its distinctive ecological niche and phylogenetic history. In other respects, however, living things are strikingly similar. They share fundamental features of structure and function that reflect their common ancestry and biophysical constraints on subsequent evolution and diversification. Some of the shared traits are at molecular and cellular levels of organization, where they reflect the biochemistry of inheritance and metabolism. Other shared traits are at higher levels of organization, where they reflect general patterns and processes of physiological performance, population dynamics, ecosystem organization and evolutionary diversification.

This duality – uniqueness and universality – of life is an outcome of evolution by natural selection. Within a population, variation among genes and individuals results in differential survival and reproduction and leads to descent with adaptive modification. Fitness is a central concept in evolutionary biology: fitter individuals have higher survival or reproduction, passing on their heritable traits to offspring. Ever since Darwin, theoretical biologists have struggled to define fitness and empirical biologists have struggled to measure it. Indeed, biologists have defined fitness in many different ways depending on the unit of measurement and level of analysis (Box 1), but at root, it is the capacity to leave descendant individuals, genes and/or quantitative heritable traits in the next generation. One corollary is that although fitness varies among individuals within a population, it is nearly equal across species in ecological assemblages. This equal fitness paradigm (EFP) is a consequence of competition for the limited supply of usable energy in the biosphere (Boltzmann 1886; Lotka 1922; Van Valen 1977, 1980) and constraints and tradeoffs in how it is allocated to production and survival (Burger et al. Brown et al 2018; 2019a). The resultant near equal fitness of species is a necessary condition for the origin and persistence of biodiversity.

For decades’ synthesis across levels of biological organization and subdisciplines of ecology and evolution were inhibited because they used different currencies (e.g., Brown 1981, 1995; but see Hutchinson 1959, 1965; Whittaker 1975; Peters 1983; Yodzis and Innes 1992; Cohen et al, 1993; De Angelis 1995). Physiologists at the individual-organism level and ecologists at the ecosystem level focused on metabolism and used biophysical 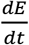 currencies: fluxes and stores of energy and materials. Ecologists at population and community levels focused on population dynamics and used numerical 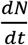 currencies: numbers of individuals. Synthesis was inhibited because it was far from straightforward to translate the biophysical data and theories of physiological and ecosystem ecology into the numerical currencies of demography and population and community ecology – and vice versa. Recently, however, ecologists have begun to focus on energy metabolism to synthesize across subdisciplines and levels of organization by linking the performance of individuals to emergent consequences for niche relations, ecosystem organization and biodiversity. The EFP is the inevitable outcome of competition and coevolution among species for the incident solar energy captured in biomass.

### An energetic definition of fitness

Energy is the ultimate limiting resource for living things. As Boltzmann (1886) perceptively wrote: “The “struggle for existence” of living beings is … for the possession of the free energy obtained, chiefly by means of the green plant, from the transfer of radiant energy from the hot sun to the cold earth.” The solar energy captured and converted into the organic molecules by photosynthesis in green plants supplies essentially all energy used by organisms. The energy in the molecules is released and used to fuel the work of living in the processes of metabolism: assimilation, respiration and production. Assimilation is the uptake of energy from the environment. Autotrophic plants take up solar energy, carbon dioxide and oxygen and capture the energy in organic molecules. Heterotrophic animals and microbes acquire biomass energy by consuming plants. Respiration is the biochemical transformation of biomass energy to fuel work: organic molecules are combined with oxygen, reduced to carbon dioxide and water, and energy is captured in ATP molecules. ATP is transported around the body, where it is converted to ADP, releasing energy to perform the work of digestion, excretion, activity, growth and reproduction. Some assimilated biomass energy is not catabolized in respiration but allocated to production and passed on to the next generation in the form of offspring growth and parental investment.

Energy is arguably the most fundamental currency of biological fitness, because the traditional components of fitness – survival, growth and reproduction – are all governed by the physical laws of energy and mass balance. Energy balance requires that each generation allocates the energy assimilated from the environment between respiration and production; the energy allocated to production is partitioned between growth and reproduction. The gain of energy due to production, is thus matched by loss of energy to mortality (see below and Fig.1). One consequence of energy balance is that fitness can be expressed in terms of energy

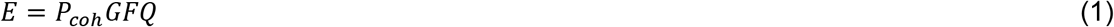

where *E* is energetic fitness in kilojoules per gram per generation, *P_coh_*, is the mass-specific rate of biomass energy production (growth plus reproduction) of the cohort of offspring produced by a parent in a lifetime in grams per gram per year, *G* is generation time in years, *Q* is the energy density of biomass in kilojoules per gram, and *F* is the fraction of lifetime biomass production that is passed through to ***surviving*** offspring in the next generation (Brown et al. 2018). So eq 1 defines the components of traditional Darwinian fitness – survival, growth and reproduction – in terms of the physical currencies and metabolic processes of energy metabolism.

**Fig. 1.**
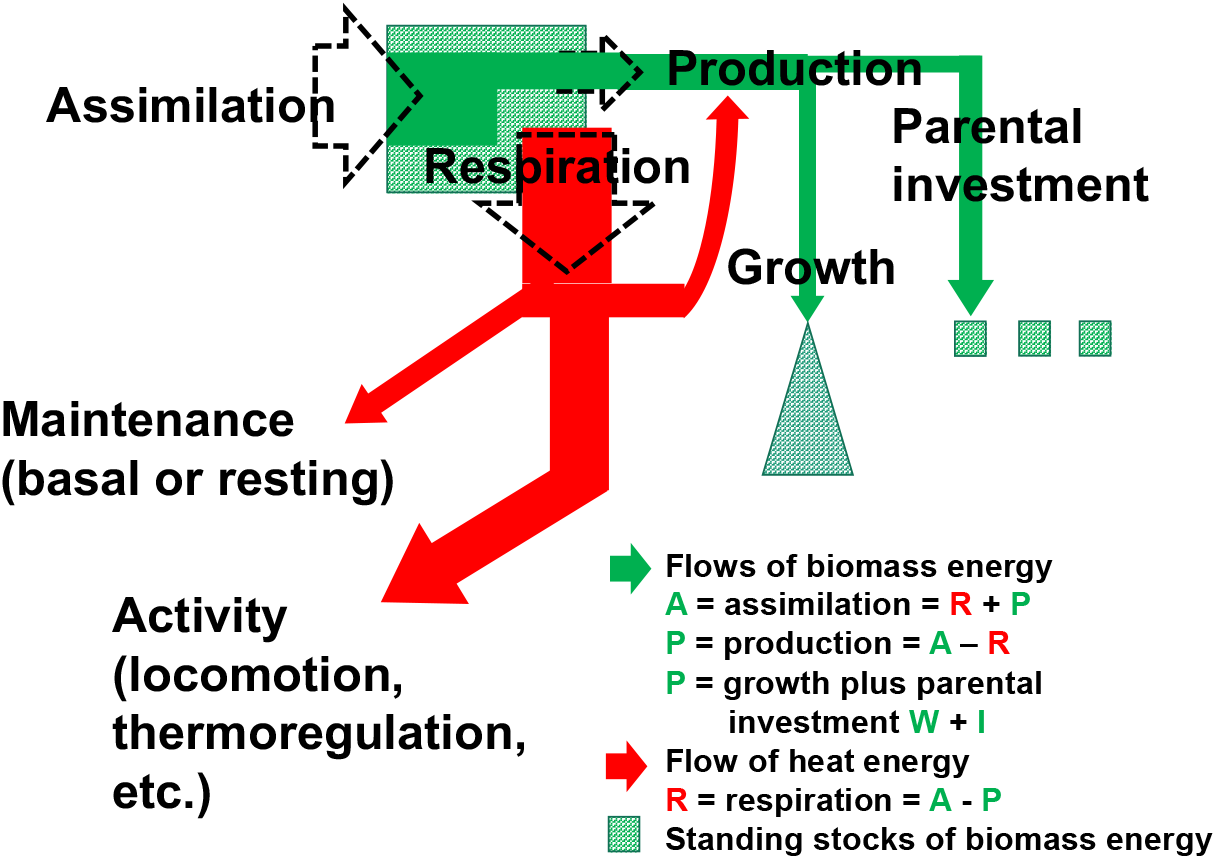
Energy balance of an individual animal. Energy assimilated from food is allocated between respiration and production; energy captured in ATP by respiration is allocated between basal or resting metabolism and activity metabolism (which includes digestion, thermoregulation, growth, immune response to pathogens, and locomotor, feeding, territorial defense, courtship and mating behaviors). Biomass energy passed to the next generation as production is allocated between offspring growth and parental investment in gametes and nutrition. Over the lifespan of an average individual at steady state, energy taken up from the environment in assimilation is returned to the environment in mortality and heat as metabolism.

This relation can be simplified by recognizing that the energy density of biomass is nearly constant across all living things: *Q* ≈ 22.4 kJ/g dry weight (see below). Consequently, eq 1 can be rewritten in terms of mass balance

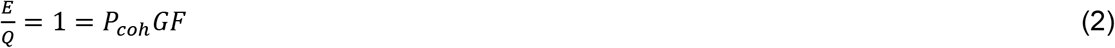

where 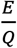 is fitness in grams per gram per generation.

This biophysical characterization of fitness based on energy and mass balance (Fig 1 and seminal eqs 1 and 2) applies to all levels of biological organization from alleles, quantitative traits and individuals within populations to species within communities, ecosystems and the biosphere. Most traditional treatments of fitness focus on the population level where variation among alleles, quantitative heritable traits, or individuals results in differential survival and reproduction (Box 1). Such variation is the basis for evolution by natural selection. Individuals with higher than average fitness have a higher probability of survival and reproduction and leave more descendants in future populations, so heritable traits that confer such advantages increase in frequency. This is consistent with energetic fitness; any differential survival (generation time) or reproduction (production rate) is necessarily reflected in a change in energetic fitness. When natural selection is operating – i.e., when a gene, trait or individual has higher than average fitness – energetic fitness is greater than 1 and increases or decreases, respectively, in the next generation. Expressing fitness in units of energy or mass instead of alleles, heritable traits or individuals and relating traditional biological currencies to their biophysical underpinnings is consistent with our understanding of fitness, natural selection and evolution at the population level. It leads, however, to new insights and perspectives in ecology and evolution at the levels of communities, ecosystems, and the biosphere.

#### Box 1. Natural selection and alternative definitions of fitness

Ever since Darwin’s theory of evolution by natural selection, biologists have struggled to define fitness. Most definitions assume that fitness is the quantity that is maximized or optimized by natural selection. Natural selection operates on variation among genes, quantitative traits and individuals **within** a single species population. When natural selection is operating, there is a departure from steady state. Fitter individuals leave more descendent heritable traits and descendants in the next generation.

Several measures of fitness are well-established in the literature of life history, evolutionary and physiological ecology.

##### 1) Rate of increase

One is based on the premise that natural selection tends to maximize the rate of increase in heritable traits that enhance reproduction or survival, and of individuals possessing such traits. Two measures are commonly used: i) the population growth rate, *r* = 1/*N dN/dt*, where *N* is the number of copies or individuals; and ii) the net reproductive rate, *R*_0_ = ∑*l_x_f_x_*, where *l_x_* is survival and *f_x_* is fecundity as a function of age *x*) (e.g., Charlesworth 1973, 1994; Charnov and Schaffer 1973; Stearns 1977, 1992; Charnov 1991, 1993; Sibly 1991, 2002; Brown and Sibly 2006). These are typically applied to quantify rates of change per unit clock time (*r*) or per generation (*R*_0) in the frequency of an allele, quantitative trait, genotype or phenotype within a population during departures from steady state when natural selection is operating.

##### 2) Maximum power

An alternative suggestion is that selection tends to maximize metabolic or reproductive power. This has been called the maximum power principle (MPP) and attributed primarily to Alfred Lotka and H.T. Odum (Lotka 1922, Odum and Pinkerton 1955, Odum 1971; see also Brown et al 1993; Hall 1995). Power is measured in units of energy per time (e.g., J/s or watts). Biological power, expressed as rate of respiration or production, increases with body mass, scaling ∝ *m*^3/4^. But larger more powerful animals are not necessarily fitter.

##### 3) Resource use efficiency

Physiological ecologists have frequently used resource use efficiency (RUE) – of carbon, water, a nutrient or some other limiting resource – as a measure of performance, with the implication that natural selection tends to maximize RUE (DeLucia and Schlesinger 1991; Chapin et al. 1997; Sterner and Elser 2002; Funk and Vitousek 2007; Hodapp et al. 2019; Vitousek 1982). Efficiency is the unitless ratio: *REU = output/input*, measured for example, in g/g or kJ/kJ. However, greater REU does not mean that more efficient organisms are necessarily fitter.

##### 4) Lifetime reproductive effort

Our energetic fitness, *E*, has similarities to the lifetime reproductive effort, LRE, of Charnov (1991, 1993; Charnov et al. 2007), in that both *E* and LRE are predicted to be approximately constant: independent of body size, fecundity, lifespan and other life history traits. Charnov’s model uses the net reproductive rate, *R*_0, as the measure of fitness and makes assumptions about how component variables scale with body mass. The big difference between LRE and *E* is that the former does not explicitly incorporate growth and pre-reproductive mortality of offspring (*W_coh_*, eg 7). Consequently, the prediction of nearly constant LRE seems to hold only for animals such as mammals and lizards (Charnov and Ernest 2006; Warne and Charnov 2008), which produce a few relatively large offspring with relatively little growth and mortality after independence. The EFP can be viewed as a more general theory that subsumes Charnov’s model of constant lifetime reproductive effort as a special case.

The above definitions differ from the energetic fitness, *E*, of the EFP. The EFP was developed by Brown et al. (2018) to account for the nearly constant value of *E* ≈ 22.4 kJ/g/generation (eqs 1 and 2) across species in different taxa, functional groups and environments. The EFP, like the Hardy-Weinburg equilibrium, uses the assumption of steady state to establish a baseline from which to quantify departures. Therefore, the EFP is distinct from but consistent with definitions of fitness that focus on evolution of traits due to natural selection. The EFP does not contradict the proposition that natural selection can act to increase the above measures of fitness (*r, R*_0, metabolic power, RUE or *L*), but only during departures from steady state when the more or less fecund, long-lived, powerful or efficient variant temporarily has higher energetic fitness.

### The equal fitness paradigm (EFP)

Deeper insights into the biophysical underpinnings of ecology and evolution come from invoking the assumption of population steady state and extending the concept of energetic fitness to multiple levels of biological organization. This leads to the equal fitness paradigm (EFP: Brown et al. 2018, Burger et al. 2019a). The steady state assumption is critical but also realistic. At steady state parents replace themselves with an equal energy content, mass and number of offspring each generation, so birth rates equal death rates, populations do not increase or decrease, species coexist and biodiversity is maintained. Eqs 1 and 2 show that species have equal energetic fitness because at steady state parents pass on 22.4 kJ/g of energy or 1 g/g of biomass to surviving offspring each generation.

In some respects, the EFP is reminiscent of the Hardy-Weinburg equilibrium (Hardy 1908; https://en.wikipedia.org/wiki/Hardy%E2%80%93Weinberg_principle) in population genetics. The HWE also makes the critical assumption of steady state as a proof that the frequency of genes and genotypes in a population do not inherently change over time. When first proposed, the HWE seemed counterintuitive to many geneticists, who assumed that the frequencies of dominant genes would increase over generations. The HWE is also tautological because it states, that at steady state, the frequencies of genes do not change. The HWE is more than just a tautology, however. It plays a seminal role in population genetics and evolution, because it provides the theoretical basis for quantifying the frequency of genes and genotypes in populations and for predicting changes in these frequencies due to natural selection, genetic drift and migration when the steady state assumption does not hold.

The EFP may also seem tautological, counterintuitive or both to biologists, who traditionally have associated fitness with variation in survival and reproduction. Indeed, components of fitness such as generation time and fecundity differ across species from microbes to mammals by many orders of magnitude. Energetic fitness is constant across species because there is a tradeoff between production rate, *P_coh_*, and generation time, *G*, such that in each generation at steady state, parents on average replace themselves with offspring containing an equal quantity of biomass and 22.4 kJ/g of energy. The product, *P_coh_GF*, is constant, even though the three variables vary across species by many orders of magnitude. The enormously varied patterns in life history strategies reflect a universal tradeoff in among the pathways of energy allocation.

Much of the variation is related to body size and temperature. The standard metabolic theory of ecology (Brown et al. 2004) predicts the scalings of *P_coh_* and *G*:

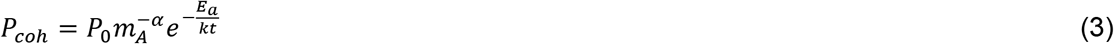

and

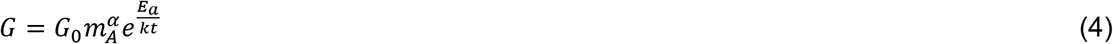

where above *P_coh_* is the mass-specific rate of cohort biomass production in grams per gram per year, *P*_0_ is a normalization coefficient, *G* is generation time in years, *m_A_* is adult body mass, *α* is the mass-scaling exponent, *e is* the root of the natural logarithm, *E_a_* is an “activation energy” which gives the temperature dependence, *k* is Boltzmann’s constant, and *t* is the temperature in Kelvin. These scaling relations apply both over ontogeny within species and across species. Standard metabolic theory predicts that *α* = 1/4 and *E_a_* ≈ 0.65 eV, equivalent to a Q10 of approximately 2.5 (Brown et al. 2004; Sibly et al. 2012a).

These equal-but-opposite scalings of production rate and generation time with body mass are well supported empirically (e.g., by Hatton et al. 2019). Their values of the exponents, −0.26 and 0.24 for what they called “growth rate” and “lifespan” respectively, are very similar to the predicted values of −1/4 and 1/4 for what we call “production rate” and “generation time”. Consequently, the product, their “lifetime growth” or our “energetic fitness” is nearly constant, ≈ 0 and independent of body size (and also temperature: not shown here, see Gillooly et al. 2001; Brown et al. 2004, 2018; Hatton et al. 2019; Burger et al. 2019a).

There is, however, considerable variation. Some of this due to lack of standardization and measurement error. Testing the EFP (eqn 1) requires consistent definitions and standardization of parameters (Box 2). Even when standardization and assumptions are met, however, there is still biologically relevant variation remaining in the four parameters of the EFP. Equation (1) provides a quantitative framework to unravel this variation. First, the relative constancy of *E* and, *Q* is striking given the diversity of life. Second, traditional metabolic scaling theory predicts most of the variation in *P_coh_* and *G* with body size and temperature as (Brown et al. 2004; Sibly et al. 2012a). Variation across taxa and functional groups generally offset, resulting in relatively invariant scaling exponents and normalization constants. Third, there is additional systematic variation in *Q* and *F*, which is not accounted for by body size and temperature. Some of this is apparently due to variation in fecundity and parental care (see below). Fourth, variation in heritable traits within a population is subject to natural selection and niche evolution. It can lead to temporal and spatial variation when the steady state assumption does not hold. We elaborate in the following sections after summarizing a metabolic theory of life history (Burger et al. 2019a).

#### Box 2: Definition and standardization of metabolic variables

The sections on energy balance and life tables highlight the importance of carefully defining and measuring the relevant variables. This has often not been done in past studies of biological scaling, metabolic ecology and in compiling the electronic databases that have been used in empirical analyses. Failure to do so can result in both systematic deviations and random errors. Relevant issues include:

i. Assimilation: We define assimilation rate as the rate of uptake of energy or matter from the environment: i.e., absorption from the gut or across the body surface. In complex metazoan animals, assimilation is ingestion of food minus excretion of feces – an important clarification, because assimilation is often ***assumed*** to be some constant fraction of gross food consumption.
ii. Respiration: If not rigorously defined and carefully measured, variation in reported respiration rates can reflect the influence of uncontrolled factors, including the biochemical composition of biomass, the oxidation pathway (e.g., aerobic vs anaerobic metabolism), level of activity (e.g., basal, resting, maximal or field metabolic rates), and other states (e.g., thermoregulation, stress or reproduction). For example, maximal respiration rates during physical activity (VO2max) can vary several fold, both across species depending on “athleticism” (e.g., sedentary sloth vs athletic spider monkey) and within individuals depending on intensity and duration of activity (e.g., 100 m dash vs 100 km ultramarathon). Reported maximal and field respiration rates are often simply ***assumed*** to be some constant multiple of basal or resting rates.
iii. Production: Reported “production rates” in the literature and electronic databases have often been measured inconsistently, i.e., as either growth or parental investment rather than both components (e.g., Peters 1983; Ernest et al. 2005; Sibly and Brown 2007; Brown et al 2018; Hatton et al. 2019).
iv. Lifespan and generation time: Several different life history times are often reported in the literature and comparative databases for specific taxa: average (often not rigorously defined), total (from birth to death), reproductive (from first to last breeding), and reflecting closer to maximum lifespans from birth to death for both wild and captive animals (e.g., Myhrvold et al. 2015; De Magalhães and Costa 2009). These are not directly comparable to each other or to generation time, so using them indiscriminately can lead to confusion and serious errors. For example, it is often ***assumed*** that generation time is the reciprocal of lifespan (*G* = 1/*t_life_*), but this is only true when *t_life_* is the average time from birth to replacement reproduction, which requires a life table for accurate determination.

### Metabolism and life history

The concept of energetic fitness integrates physics and biology, metabolism and life history. It addresses how the energetic performance of individual organisms affects their contributions to future generations, interactions with other species and the abiotic environment and coevolutionary dynamics. The life history is the schedule of survival and reproduction over a lifetime; it determines the contributions of genes, traits and individuals to fitness in the next generation. It is subject to the physical laws of energy and mass balance and the biological laws of demography and population dynamics.

#### 1) Energy balance of an individual

All organisms obey the physical laws of energy and mass balance (Lotka 1922; Odum 1971; Kooijman 2000). Figure 1 diagrams the energy balance of an individual animal in terms of the uptake, transformation and expenditure of metabolic energy.

##### i) Assimilation

Animal metabolism is fueled by consumption and assimilation of food. As Boltzmann pointed out (see above), the energy used by organisms comes ultimately from the sun and fixed in the organic molecules by photosynthesis in plants. Animals take up biomass energy from the environment by consuming and assimilating plants or other food from animals (and from solar energy in plants). The energy acquired from assimilation is processed and transformed within the body and allocated between respiration and production.

##### ii) Respiration

Most of the energy acquired by assimilation is expended on respiration. The organic molecules in food are eventually broken by oxidative metabolism to yield the waste products carbon dioxide, water and heat, which are released back into the environment. Usable energy is temporarily stored in ATP, which is subsequently metabolized to perform the physical work of living: maintenance, growth and activity. Respiration is what Kleiber (1932) and subsequent physiologists refer to as “metabolism” or “metabolic rate”.

Animal physiologists often use standardized conditions in the laboratory to measure different levels of metabolic performance: i) basal or resting metabolic rates (BMR or RMR), the respiration rate of an inactive, fasting individual in its thermoneutral zone (growing animals and those engaged in thermoregulation, locomotion, feeding, territorial defense, courtship, reproduction and immune response to pathogens have higher rates); ii) maximal metabolic rates (VO2 max), elevated respiration rates during activity or stress); and iii) field metabolic rate (FMR), average respiration rates of free-living animals in nature. All of these levels of respiratory metabolism are correlated to each other; they may differ in normalization coefficients, but scale similarly with body mass: as 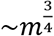 for whole-organism rates and 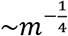 for mass-specific rates, and with temperature (e.g., Peters 1983; Calder 1984; Schmidt-Nielson 1984; McNab 2002; Speakman 1997; Gillooly et al. 2001; Savage et al. 2004; but see Nagy 1999, 2005; Glazier 2010; Speakman and Krol 2010). Since these measures of respiration do not include uptake and expenditure for growth and reproduction, they do not represent energy balance for an individual over its lifespan.

##### iii) Production

A fraction of the biomass acquired over the lifetime is not broken down in respiration but stored temporarily in the body as production of “net new biomass”. The macromolecules may be altered by metabolic biochemistry, but they are passed on with most of their energy intact and allocated between somatic growth and parental investment in reproduction.

As shown in Figure 1, mass-energy balance for an animal over its lifetime requires that input equals output, so: i) assimilation, *A_ind_*, or uptake from food; equals the sum of ii) respiration, *R_ind_*, for synthesis and expenditure of ATP plus iii) plus production, *W_ind_*, for growth and reproduction:

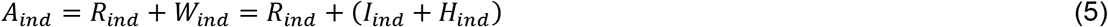

where all rates are for whole-organism per generation. Production has two components: *H_ind_*, biomass accumulated in the body as ontogenetic growth, and *I_ind_*, parental investment in reproduction in the form of gametes and post-fertilization nutrition (e.g., pregnancy, lactation and feeding in mammals).

#### 2) Demography and life tables

All organisms are mortal. Life persists because parents replace themselves with offspring in succeeding generations. The life history is the schedule of survival, growth and reproduction over an entire life cycle of one generation. Animals exhibit a spectacular diversity of life histories including sexual and asexual reproduction, semelparity and iteroparity, determinate and indeterminate growth, and orders-of-magnitude variation in body size of parent and offspring, fecundity and generation time. This variation notwithstanding, however, the biological law of demography dictates that in sexual species at steady state, two parents replace themselves over their lifetimes with two offspring; regardless of the number of offspring born, which can vary from a few to billions, only two survive to reproduce and complete the life cycle.

##### i) Net reproductive rate

Fundamental features of demography are typically presented in a life table, which quantifies survival, *l_x_*, and fecundity, *f_x_*, as functions of age, *x*. The net reproductive rate, *R*_0_ = ∑*l_x_f_x_*, is a commonly used measure of fitness (Box 1). At steady state, fitness is neutral, the population does not grow because the birth rate equals the death rate, *R*_0_ = 1, the fecundity of the average female is 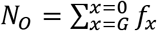, and the number of surviving offspring, *N*, decreases with age from *N*_0_ at *x* = 0 to 2 at *x* = *G*. There are hundreds of published life tables for animal species with diverse life histories.

##### ii) Metabolic life table

A complete accounting of allocation of energy or mass to fitness over an entire life cycle requires additional data on ontogenetic growth: i.e., body mass or energy content of individuals of age *x*. Then it is possible to construct a metabolic life table (MLT; Brett 1983; Burger et al. 2020). Unfortunately, sufficiently complete and detailed data to construct an MLT are available for only a few well-studied species. Furthermore, such information is not available in large comparative databases such as FishBase (Froese and Pauly 2019), AMNIOTE (Myhrvold et al. 2015), AnAGE (De Magalhães and Costa 2009), COMADRE (Salguero-Gómez et al. 2016) or DEB (Marques et al. 2018) due to lack of standardization (Box 2) and steady-state assumptions. In Table 1 and Fig. 4 we present data on energetics and demography for eight animal species with diverse life histories. Note that despite wide variation in the underlying parameters, at steady state all of them obey the laws of energy-mass balance and demography, and they have equal fitness with *E* = 22.4 kJ/g/generation.

**Fig. 2.**
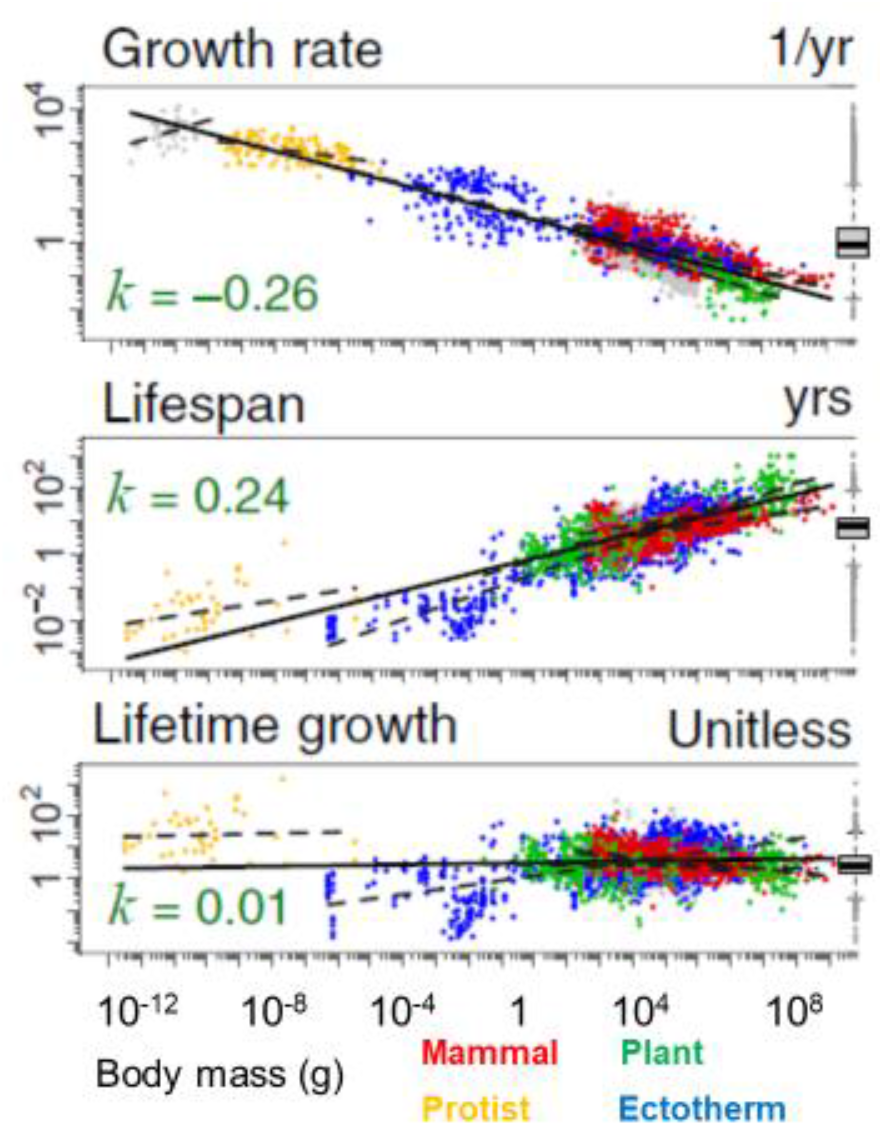
Data from Hatton et al. (2019) showing the scalings of individual production rate, generation time, and individual lifetime production as a function of body mass for four major taxonomic/functional groups plotted on logarithmic axes. Note the close agreement between the fitted exponents and the predicted values: −0.26 vs −0.25, 0.24 vs 0.25 and 0.01 vs 0.00, respectively. Note gray points are bacteria.

**Fig. 3.**
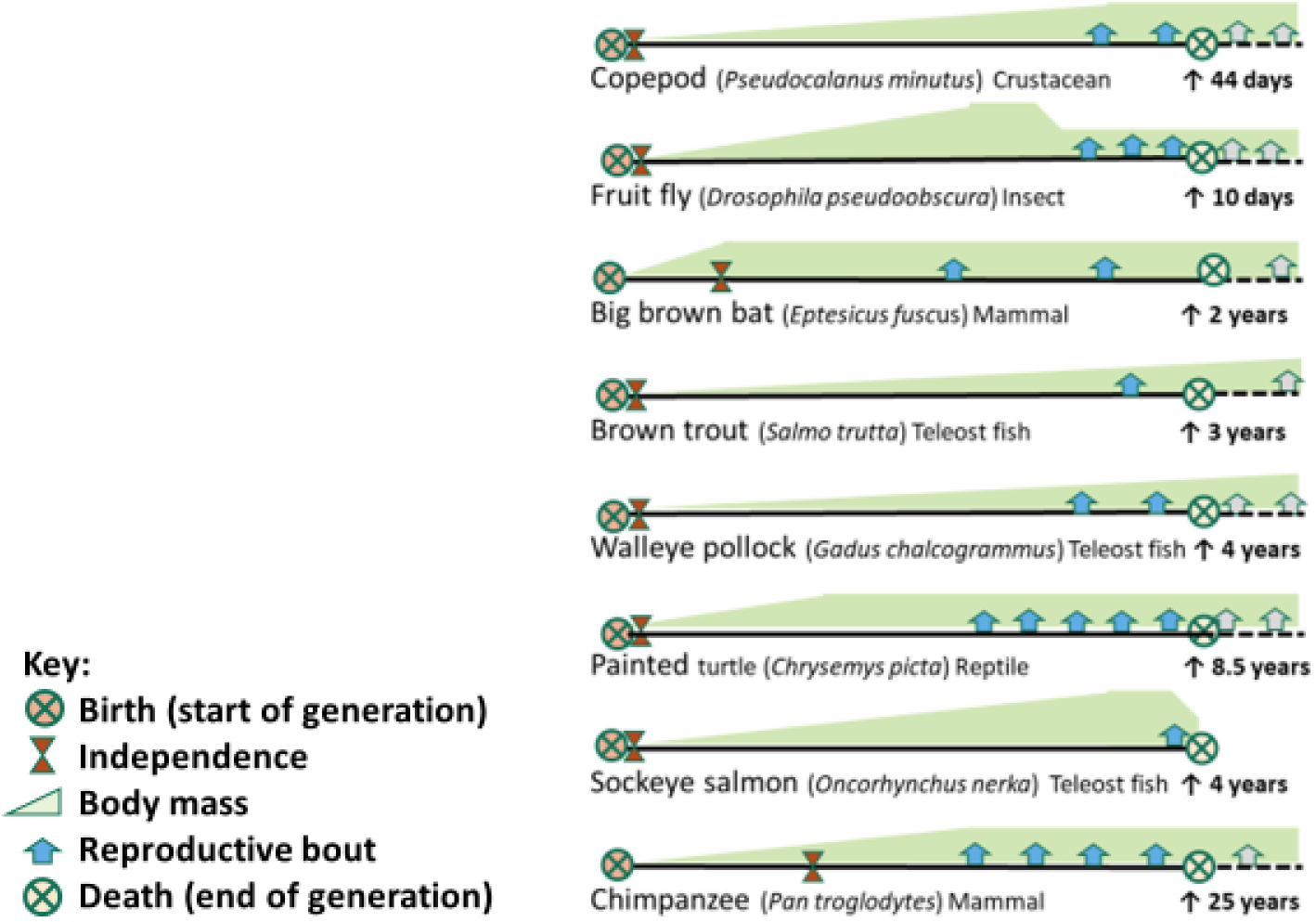
Life history schedules for various animal species, showing the relation of generation time (solid line between ⊗-birth and ⊗-death value given by ↑) to times of other events. We define generation time as time from birth to death of an average parent that replaces itself by leaving two surviving offspring. Note sockeye salmon is the only semelparous species, which dies after a single reproductive bout. In contrast in iteroparous species, some individuals live longer, continue to reproduce and sometimes grow beyond the average generation time defined here. Note that other life history times given in the literature and electronic databases, such as average, maximum and reproductive lifespans and age to first reproduction, are not the same as generation time. See also text on definitions and standardizations.

**Fig. 4:**
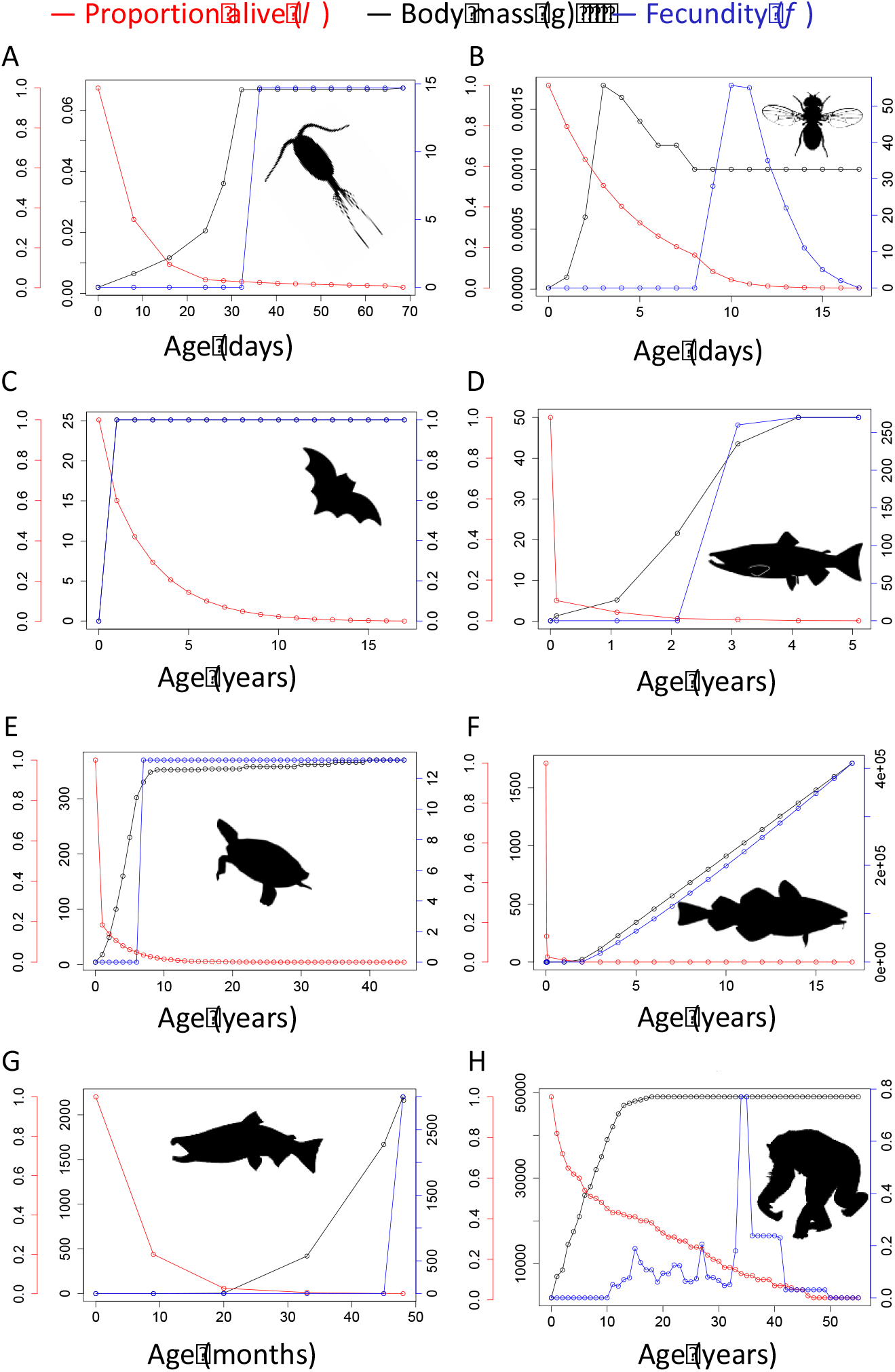
Metabolic life histories of eight animal species chosen based on data availability and to represent a diversity of taxa, body sizes and lifestyles. Included here are values for teleost fish including sockeye salmon, which grow for four years and then have just one bout of reproduction (data from Brett 1983; Brett 1986; Furnell and Brett 1986; Larkin McDonald 1968); and marine walleye pollock (data from Hinckley 1987; Houde 1989; Houde 1997 [used for larval stages]; Houde and Zastrow 1993; Smith 1979) and fresh-water brown trout (data from Brown 1946; Horton 1961; Dartmoor wild trout project 2016), which continue to grow and reproduce throughout life; mammals such as the big brown bat (data from O’Shea et al. 2010; O’Shea et al. 2011) and chimpanzee (data from Hill et al. 2001; Bronikowski et al. 2016), which provide nutrition and parental care to produce a few large offspring; a reptile, painted turtle (data from Wilber 1975a,b), which is relatively long-lived and produces a clutch of eggs each year; and two very small invertebrates a marine copepod (data from McLaren 1974; Frost 1989; Hunley and Boyd 1984) and terrestrial fruit fly (data from Church and Robertson 1966; Rosewell and Shorrocks 1987; Roberston and Sang 1944; Smith 1958; Tantawy and Vetukiv 1960), which both lay miniscule eggs but have very different patterns of growth and mortality. Life histories of eight species, depicted as trajectories over the lifespan for: growth (body mass), survival (number of offspring alive), and fecundity (number of offspring per female). Note the scales of the axes, which indicate the magnitude of variation within and between species. (See supplemental data). Icons reused from phylopic.org under https://creativecommons.org/licenses/by/3.0/ with following credits: Chimpanzee T. Michael Keesey (vectorization) and Tony Hisgett (photography). All other icons reused under Public Domain.

**Table 1:**
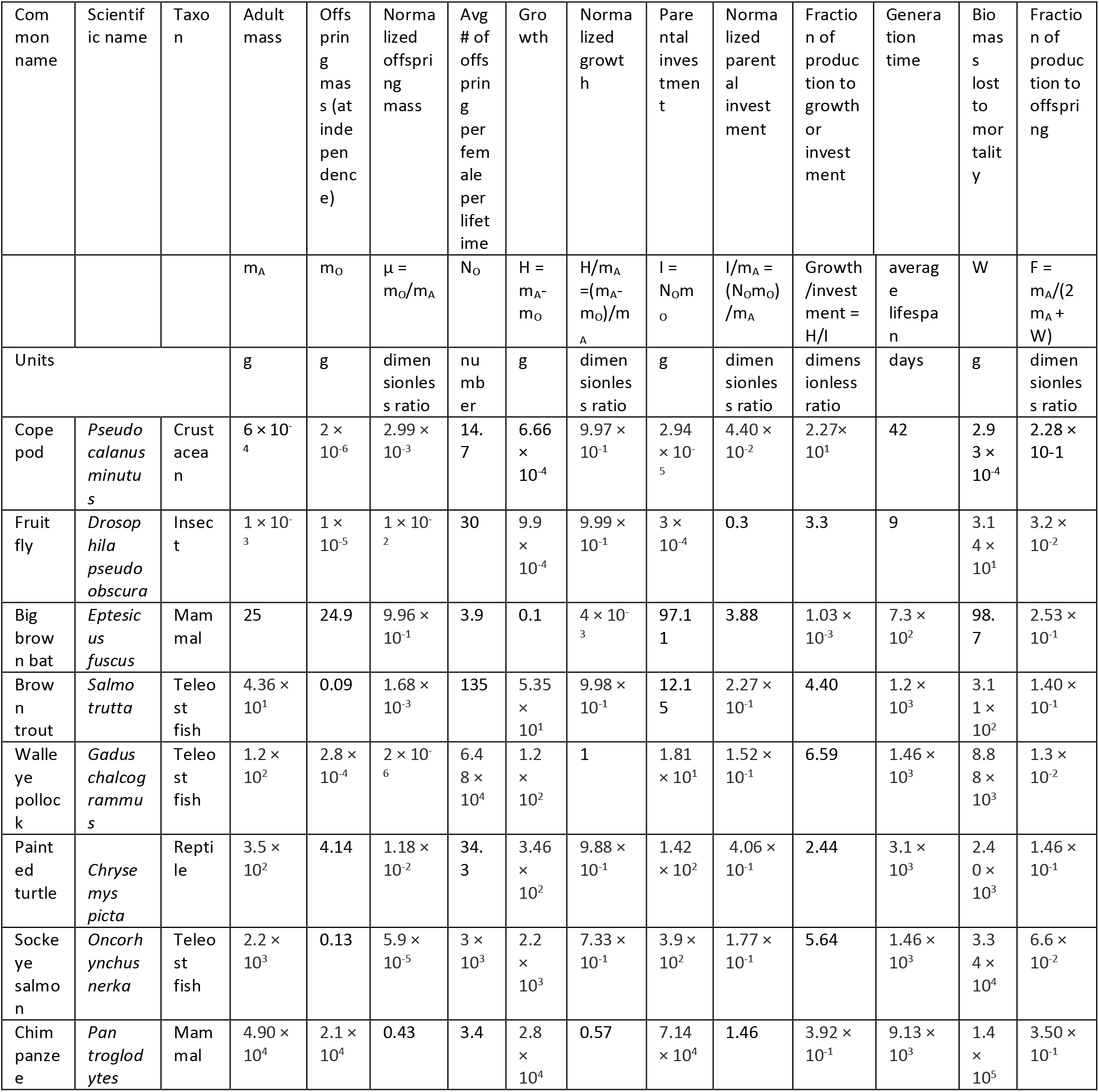
Values of life history parameters for the animal species illustrated in Figure 3. All traits vary by more than two orders of magnitude across the eight species, but the variation is constrained. Large teleost fish produce thousands of miniscule larvae that suffer high mortality as they grow to maturity. Tiny invertebrates (copepod and fruit fly) produce smaller numbers of comparably-sized larvae that suffer lower mortality because of shorter generation times. Mammals (bat and chimpanzee) invest extensive nutrition and care to produce a few relatively large offspring which suffer relatively low mortality.

| Common name | Scientific name | Taxon | Adult mass | Offspring masses (at independence) | Normalized offspring mass | Avg # of offspring per female per lifetime | Growth | Normalized growth | Parental investment | Normalized parental investment | Fraction of production to growth or investment | Generation time | Biomass lost to mortality | Fraction of production to offspring |
| --- | --- | --- | --- | --- | --- | --- | --- | --- | --- | --- | --- | --- | --- | --- |
| | | | $m_A$ | $m_O$ | $\mu = m_O/m_A$ | $N_O$ | $H = m_A - m_O$ | $H/m_A = (m_A - m_O)/m_A$ | $I = N_O m_O$ | $I/m_A = (N_O m_O)/m_A$ | Growth/investment = $H/I$ | average lifespan | $W$ | $F = m_A/(2m_A + W)$ |
| Units |  |  | g | g | dimensionless ratio | number | g | dimensionless ratio | g | dimensionless ratio | dimensionless ratio | days | g | dimensionless ratio |
| Copepod | *Pseudocalanus minutus* | Crustacean | $6 \times 10^{-4}$ | $2 \times 10^{-6}$ | $2.99 \times 10^{-3}$ | 14.7 | $6.66 \times 10^{-4}$ | $9.97 \times 10^{-1}$ | $2.94 \times 10^{-5}$ | $4.40 \times 10^{-2}$ | $2.27 \times 10^1$ | 42 | $2.93 \times 10^{-4}$ | $2.28 \times 10^{-1}$ |
| Fruit fly | *Drosophila pseudoobscura* | Insect | $1 \times 10^{-3}$ | $1 \times 10^{-5}$ | $1 \times 10^{-2}$ | 30 | $9.9 \times 10^{-4}$ | $9.99 \times 10^{-1}$ | $3 \times 10^{-4}$ | 0.3 | 3.3 | 9 | $3.14 \times 10^1$ | $3.2 \times 10^{-2}$ |
| Big brown bat | *Eptesicus fuscus* | Mammal | 25 | 24.9 | $9.96 \times 10^{-1}$ | 3.9 | 0.1 | $4 \times 10^{-3}$ | 97.11 | 3.88 | $1.03 \times 10^{-3}$ | $7.3 \times 10^2$ | 98.7 | $2.53 \times 10^{-1}$ |
| Brown trout | *Salmo trutta* | Teleost fish | $4.36 \times 10^1$ | 0.09 | $1.68 \times 10^{-3}$ | 135 | $5.35 \times 10^1$ | $9.98 \times 10^{-1}$ | 12.15 | $2.27 \times 10^{-1}$ | 4.40 | $1.2 \times 10^3$ | $3.11 \times 10^2$ | $1.40 \times 10^{-1}$ |
| Walleye pollock | *Gadus chalcogrammus* | Teleost fish | $1.2 \times 10^2$ | $2.8 \times 10^{-4}$ | $2 \times 10^{-6}$ | $6.48 \times 10^4$ | $1.2 \times 10^2$ | 1 | $1.81 \times 10^1$ | $1.52 \times 10^{-1}$ | 6.59 | $1.46 \times 10^3$ | $8.88 \times 10^3$ | $1.3 \times 10^{-2}$ |
| Painted turtle | *Chrysemys picta* | Reptile | $3.5 \times 10^2$ | 4.14 | $1.18 \times 10^{-2}$ | 34.3 | $3.46 \times 10^2$ | $9.88 \times 10^{-1}$ | $1.42 \times 10^2$ | $4.06 \times 10^{-1}$ | 2.44 | $3.1 \times 10^3$ | $2.40 \times 10^3$ | $1.46 \times 10^{-1}$ |
| Sockeye salmon | *Oncorhynchus nerka* | Teleost fish | $2.2 \times 10^3$ | 0.13 | $5.9 \times 10^{-5}$ | $3 \times 10^3$ | $2.2 \times 10^3$ | $7.33 \times 10^{-1}$ | $3.9 \times 10^2$ | $1.77 \times 10^{-1}$ | 5.64 | $1.46 \times 10^3$ | $3.34 \times 10^4$ | $6.6 \times 10^{-2}$ |
| Chimpanzee | *Pan troglodytes* | Mammal | $4.90 \times 10^4$ | $2.1 \times 10^4$ | 0.43 | 3.4 | $2.8 \times 10^4$ | 0.57 | $7.14 \times 10^4$ | 1.46 | $3.92 \times 10^{-1}$ | $9.13 \times 10^3$ | $1.4 \times 10^5$ | $3.50 \times 10^{-1}$ |

###### Box 3: Scaling relations

For centuries, natural scientists have been intrigued by the variation in forms and functions of animals. With the advent of more modern methods of measurement and analysis, pioneers such as D’Arcy Thompson (1917), Huxley (1932), Kleiber (1932), and Brody (1945) and successors such as McMahon and Bonner (1983), Peters (1983), Calder (1984), and (Schmidt-Nielsen 1984) delved into these relationships, showing that relatively simple equations can describe how diverse anatomical, physiological, behavioral and ecological traits scale with body size and temperature. In this paper, we have followed standard metabolic theory used in the scaling relations in eqs 3 and 4 (above). We recognize that throughout the history of biological scaling there has been debate about: i) the theoretically predicted or empirically estimated values of the exponents and normalization coefficients and ii) the magnitudes and sources of the remaining variation.

i. Scaling exponents: There is a longstanding debate about whether in eqs 3 and 4 the scaling exponent, *α* = 1/4 or 1/3. Ever since Kleiber’s (1932), most investigators have favored the quarter-power even though the mechanistic basis remained obscure (e.g., Peters 1983; McMahnon and Bonner 1983; Calder 1984; Schmidt-Nielsen 1984; Brown et al. 2004; Sibly et al. 2012a). But a few advocated a geometric third powers in part because heat is dissipated from the body surface which scales as *m*^2/3^ (e.g, Rubner 1883; Heusner 1991; Glazier 2005, 2010; Speakman and Król 2010). There has also been debate whether a simple Boltzmann exponential term is adequate to characterize temperature dependence, or some more complicated expression should be used (e.g., see Gillooly et al. 2001; Knies and Kingsolver 2010).
ii. Normalization coefficients: Often there is statistically significant variation in the normalization coefficient: e.g., the values of *P*_0_ and *G*_0_ in eq 3 and 4. Sometimes this is due to lack of standardization of definitions or measurements. For example, failure to standardize level of activity (e.g., time and speed of running or swimming) can lead to misinterpretation of the apparent differences (Speakman and Król 2010). Sometimes variations in normalization coefficients are “real” differences in metabolic performance among individuals (e.g., in respiration rates between the larval, pupal and adult life history stages of holometabolous insects; e.g., Llandres et al. 2015) or between species (e.g., in production rates and lifespans between birds and mammals).
iii. Error: Failure to rigorously define terms and standardize measurements results in error variation because the data are not comparable (see box Box 3). Such variation can increase the magnitude of “unexplained” variation or introduce systematic bias (e.g., for biomedical example see Dhurandhar 2015; Allison et al. 2016). Increased use of large databases and informatics is contributing enormously to macroecology and other areas of biology. But careful standardization and accurate measurement will be required to have confidence in empirical studies, regardless of whether they are inductive patterns or formal deductive tests of hypotheses.
iv. Statistical analyses: Technological advances – in hardware, software and ‘big data’ – have been accompanied by an explosion of statistical methods to characterize the magnitudes and sources of variation and distinguish among alternative models and hypotheses. We will just point out that statistics as a sub-discipline of mathematics, and its applications to other sciences are continually changing. There is no single “correct” method or “right” answer, and the best choice will likely change in a few years. The challenge is to continue to ask new questions and to expect only better – but always imperfect – answers. The EFP synthesized herein provides a unified framework to guide these questions.

#### 3) Energy density of biomass

The EFP and its seminal equations 1 and 2 call attention to the importance of *Q*, the energy density of biomass. *Q* is a biological constant: ≈ 22.4 kJ/g dry weight (≈7 kJ/g wet weight). It varies less than 2-fold across organisms spanning 20 orders of magnitude in body mass from microbes to mammals and trees (Brown et al. 2018). The value of *Q* is determined by biochemistry, because biomass is composed of similar proportions of carbohydrates (~17 kJ/g), proteins (~17 kJ/g), and lipids (~34 kJ/g). This was recognized more than 50 years ago (e.g., Cummins and Wuychek 1967), but has gone largely unappreciated in metabolic ecology. It is another universal characteristic of life at the molecular level – and one with profound implications for biological energetics from physiology to ecosystem ecology. Importantly, the constancy of *Q* allows the EFP and life history parameters to be expressed in units of mass, which are more easily measured and readily available in the literature than units of energy.

#### 4) Production of a cohort

The EFP is explicitly couched in terms of mass-energy and demographic balance of a non-growing population or the cohort of offspring produced by an average parent at steady state (Brown et al. 2018; Burger et al. 2019a):

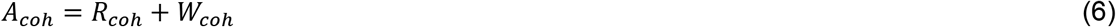

where *A_coh_* and *R_coh_* are the total assimilation and respiration of all offspring produced by a female parent, and *W_coh_* is the biomass of the entire cohort of offspring produced by the parent. So *W_coh_* is the sum of the body masses of all offspring when they die, including the two that replace the parents. So

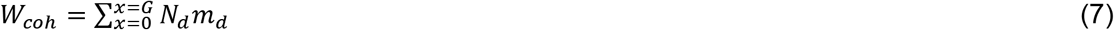

where *N_d_* is the number of offspring dying at age *x* and *m_d_* is the mass of those offspring when they die. Note that the mass-specific cohort biomass production of the EFP, 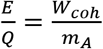 (eqs 2 and 6) is not the same as 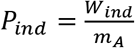 in eq 5, which has variously been called “individual production”, “maximal production”, “growth” or simply “production”. These measures do not include the production attributable to a parent in the growth of offspring that die before maturing and reproducing.

#### 5) Efficiency of production

Rarely considered in treatments of life history is the quantity *F* in eqs. 1 and 2 (but see Brett 1986). 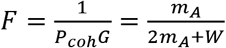 is the fraction of biomass energy produced in one generation that is incorporated into ***surviving*** offspring in the next generation, so it is a measure of efficiency of reproduction. Among the eight species in Fig. 4, values of *F* varied by more than an order of magnitude: from 0.013 to 0.032 in walleye pollock and copepod to 0.30 to 0.35 in chimpanzee and brown bat. [Note that *F* is not the same as the trophic efficiency 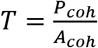 of a species in the ecosystem, another consequential mass-energy balance.]

Especially relevant to ecology is the complementary fraction (1 – *F*), the fraction of biomass produced by a cohort each generation that is lost to pre-reproductive mortality, left in the ecosystem, and mostly consumed by other organisms (predators and decomposers). Variation in *F* and (1 – *F*) is closely related to the number and relative size of offspring. For example, mammals, which have extensive parental care and produce a few relatively large offspring pass on more than half of cohort production to the next generation; they are more efficient energetically than the fish and invertebrates that produce large numbers of tiny offspring but lose more than 90% (sometimes much more than 99%) of cohort production to pre-reproductive mortality in the ecosystem.

#### 6) Toward a more complete theory

A metabolic theory of life history is still a work in progress. Recent advances highlight the promise of a synthesis based on 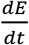 currencies and first principles of physics and biology (Burger et al. 2019a; Hatton et al. 2019; Burger et al. 2020). As exhibited in Fig. 4, animals exhibit wide variations in life histories. The EFP offers insights into the causes and consequences and the magnitudes and extents of this variation. Burger et al. (2019) showed how the basic life history tradeoffs – between generation time and production rate, offspring growth and parental investment, and number and size of offspring – are due to fundamental constraints of mass-energy balance and demography.

Other important patterns and processes remain to be explained. For example, there is necessarily a mechanistic linkage between growth and mortality, and more generally between the 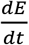 variables of metabolic ecology and the 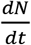 variables of demography and population dynamics. Progress will require both theoretical and empirical advances: new models of key processes and new compilations of data that address the issues of accuracy and standardization in (Box 1 and 2).

##### Box 4: Biological time and a general explanation for Kleiber’s law

The equal but opposite scaling of biological rates and times with body mass – as 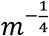 and 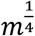, respectively (eqs 3 and 4) means that the product of rates and times is an invariant quantity: 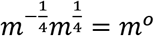. A classic example is the product of heart rate and lifespan gives the number of heartbeats in a lifetime, independent of body mass and nearly constant: ≈ 2 billion, across mammals from mice to elephants. Similarly, the EFP gives energetic fitness as the invariant product, *E* ≈ 22.4 kJ/g/generation, of production rate, *P_coh_*, times generation time, *G* (eqs 1 and 2). The central importance of generation time as a component of fitness and its equal but opposite scaling with body size and temperature (eqs 1–4) suggests that it is a universal feature of life and an alternative explanation for Kleiber’s law: metabolic rate scaling as the 3/4-power of body mass.

Ever since Kleiber’s (1932) seminal study, theoreticians have tried to explain why metabolic processes scale with quarter powers of mass or volume rather than the third powers that would be expected on the basis of Euclidean geometry. We propose that biological rates scale with quarter powers of mass because time is the fourth dimension. This is well established in theoretical physics but not generally extended to biology (but see Blum 1977; Ginzburg, and Damuth 2008). We start with the observation that life is fundamentally four-dimensional. Three are the standard dimensions of Euclidean geometry; the static characteristics of organisms’ scale as third powers of body volume: 1) length (e.g., of limbs, guts and nerves) scales as 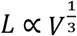; 2) surface area (e.g., of skin, lungs, gills, guts and leaves) scales as 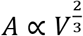; and volume or mass (e.g., of organelles and organs) scales as 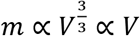. A fourth dimension, time, is necessary to capture dynamics, and characteristic times of biological processes scale as 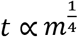. Empirical evidence indicates that biological times – spanning many orders of magnitude from the milliseconds of biochemical reactions and muscle contractions at molecular and cellular levels, to years of lifespans and population cycles at organism to ecosystem levels – scale as the 1/4-power of body mass (Fig. 5: Lindstedt and Calder 1981; Peters 1983; Calder 1984; Brown et al 2004; Andersen et al. 2016; Hatton et al. 2019).

We suggest that fitness, an existential feature of life, is fundamentally four-dimensional with fourth dimension reflecting the scaling of biological time. This leads to our hypothesis that Kleiber’s law, the *m*^3/4^ scaling of respiration rate, is the result rather than the cause of the *m*^1/4^ scaling of lifespan or generation time. Assuming that *G* ∝ *m*^1/4^ and substituting eq 4 into eq 2, we have

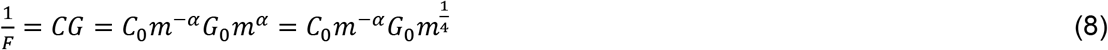

and *α* = −1/4 for mass-specific production rate. The other metabolic rates (assimilation, and resting and active respiration) scale with similar exponents (but different normalization coefficients), and whole-organism rates scale with *α* = 3/4 as in Kleiber’s law. This means that the quantity of energy expended by a gram of tissue in a lifetime is approximately invariant, independent of the size of the organism: ∝ *m*^-1/4^*m*^1/4^ ∝ *m*^0^ (e.g., Fig. 2; Andresen et al. 2002; Speakman 2005; Atanasov 2007; Ginzburg, and Damuth 2008; Brown et al. 2018; Hatton et al. 2019; Burger et al. 2019a).

The EFP raises questions about the nature of biological time. Our hypothesis (Box 4) that Kleiber’s *m*^3/4^ scaling of metabolic rate reflects the *m*^1/4^ scaling of biological time broadly and generation time in particular, is more general and widely applicable than existing models based on vascular and other systems that supply metabolites and remove wastes (e.g., West et al. 1997; West et al. 1999; Banavar et al. 2002, 2010; Aitkenhead et al. 2020). The 3/4-power scaling of respiration rate with body mass is pervasive across the animal kingdom, occurring in diverse taxonomic and functional groups that have distinctly different anatomies and physiologies for assimilating, transporting and excreting the substrates and wastes of respiratory metabolism (e.g., Peters 1983; Brown et al. 2004). These include the guts, lungs, gills, arteries, veins, tracheales, kidneys and Malpighian tubules of terrestrial, freshwater and marine species. So, our explanation subsumes most existing network models by suggesting that they are all solutions to the general problem of fueling metabolism: since time scales as *m*^1/4^, whole-organism production rate and other rates necessarily scale with quarter powers (Gillooly et al. 2001,2002; Savage et al. 2008; Banavar et al. 2010; Maino et al. 2014). This leads to a general theory for scalings of animal form and function across: i) body mass from milligrams to tonnes, ii) temperature from 0-60°C; iii) time from milliseconds to centuries, and iv) level of organization from molecules and cells to populations and ecosystems.

### The EFP and the Red Queen

The diversity of species in ecological assemblages at all scales, from local populations and communities to the global biota, is a corollary of the equal fitness paradigm: equal fitness is a necessary condition for coexistence and persistence. Within species populations, however, individuals vary in energetic fitness, *E*, and in heritable traits that determine the values of the parameters in eq 1. An exception is the parameter *Q* ≈ 22.4 kJ/g, which is nearly constant so there is virtually no heritable variation for selection to act on. There are, however, orders-of-magnitude variations in the other parameters: growth and reproduction (production rate, *P_coh_*), survival (generation time, *G*), and efficiency of production (*F*, the fraction of production passed from parent to offspring). Why doesn’t natural selection increase energetic fitness, *E*, by acting on this variation and resulting in increases in these constituent parameters?

There are at least three inter-related reasons. First, is depletion of genetic variation. Continual selection on any trait leads to reduced heritable variation and progressively slower change in fitness. Second is ecological compensation (Calow and Sibly 1983). If there is heritable variation for production rate, *P_coh_*, survival, *G*, or reproductive efficiency, *F*, natural selection will favor individuals and traits with higher values, they will increase in the population and the population will grow. But such dynamics will be transient. Ultimately population growth will be limited by some environmental factor (carrying capacity), the initial advantage will be lost, and further increases will not occur. The third reason why energetic fitness and its component traits do not increase indefinitely is because of Red Queen interactions and coevolution. As Boltzmann (1886, see above) pointed out, the biomass energy produced by photosynthesis is the ultimate limiting resource for living things. Van Valen (1973; 1980) characterized the ecological and evolutionary dynamics of species in ecological assemblages as a “zero sum game” of competition for energy. He coined the term Red Queen after the character in Lewis Carroll’s *Through the Looking Glass* who famously said “It takes all the running you can do, to keep in the same place.”

A simplified characterization of the Red Queen zero sum game is diagrammed in Fig. 6. The overall supply of usable biomass energy is set by net primary production of the ecosystem (Hutchinson 1959). There is continual selection on each species, subject to the Malthusian-Darwinian dynamic (MDD: Brown 1995; see also Nekola et al. 2013), to increase its share. The energy used by each species is set jointly by its intrinsic biological traits and local environmental conditions (e.g., Violle et al. 2012; Enquist et al. 2015; Burger et al. 2019b). Together, these define the unique ecological niche. Selection leads to the divergence of traits, such as body size, food requirement, territorial and antipredator behavior, etc., and the dispersion of niches along axes of environmental variation, such as temperature, nutrients, physical structure, predation risk, etc. (Whittaker 1970; Hall et al. 1992; Blonder et al. 2014). Moreover, as any species increases in abundance, its biomass becomes a more attractive resource for consumers: predators, parasites and pathogens. The collective result of the ecological interactions and coevolution of species is that all the biomass energy produced in the ecosystem tends to be used. At fine scales of time and space the niches and abundances are in constant flux as each species competes in the zero-sum game. At larger scales, however, there is an approximate steady state as increases in some species are balanced by decreases in others and biodiversity is maintained. At steady state the fitness of all species is very nearly equal: each is close to its carrying capacity and none exhibits more than a temporary advantage in space and time.

**Fig. 5.**
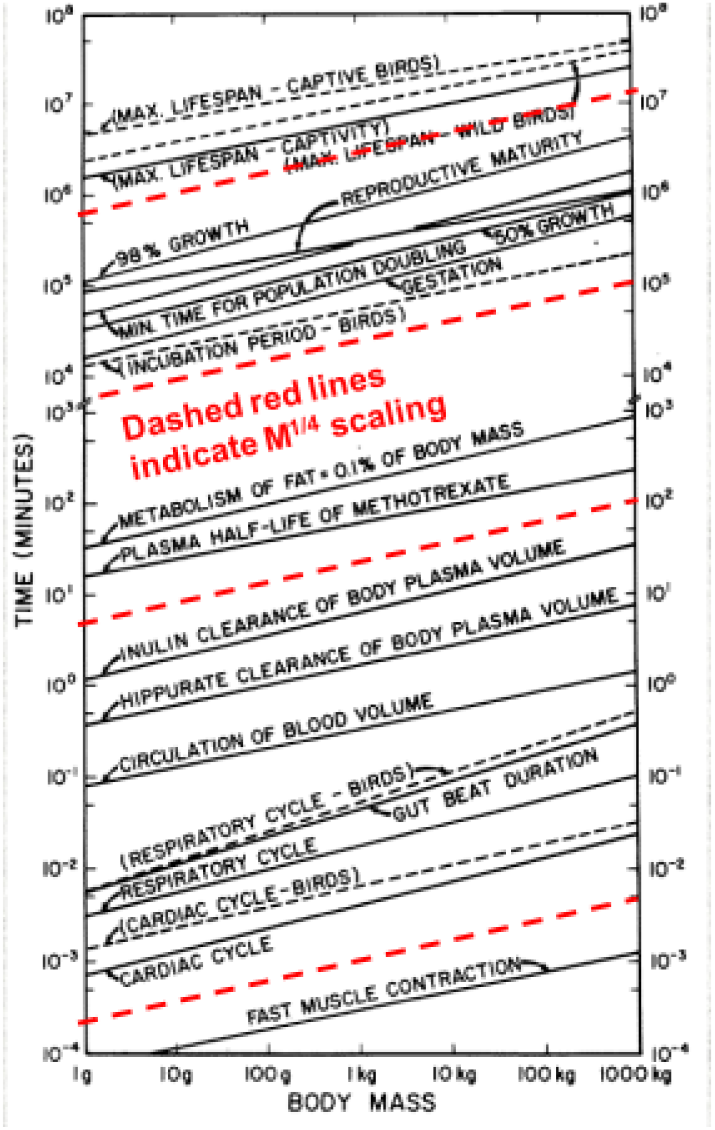
Data and analysis from Lindstedt and Calder (1981) showing that biological times scale as the 1/4 power of body mass. The authors have plotted the characteristic times of biological processes spanning 12 orders of magnitude, from milliseconds to centuries, in mammals and birds spanning 6 orders of magnitude in body mass from shrews and hummingbirds to elephants and ostriches. For reference, we have plotted dashed red lines indicating 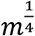 scaling.

**Fig. 6.**
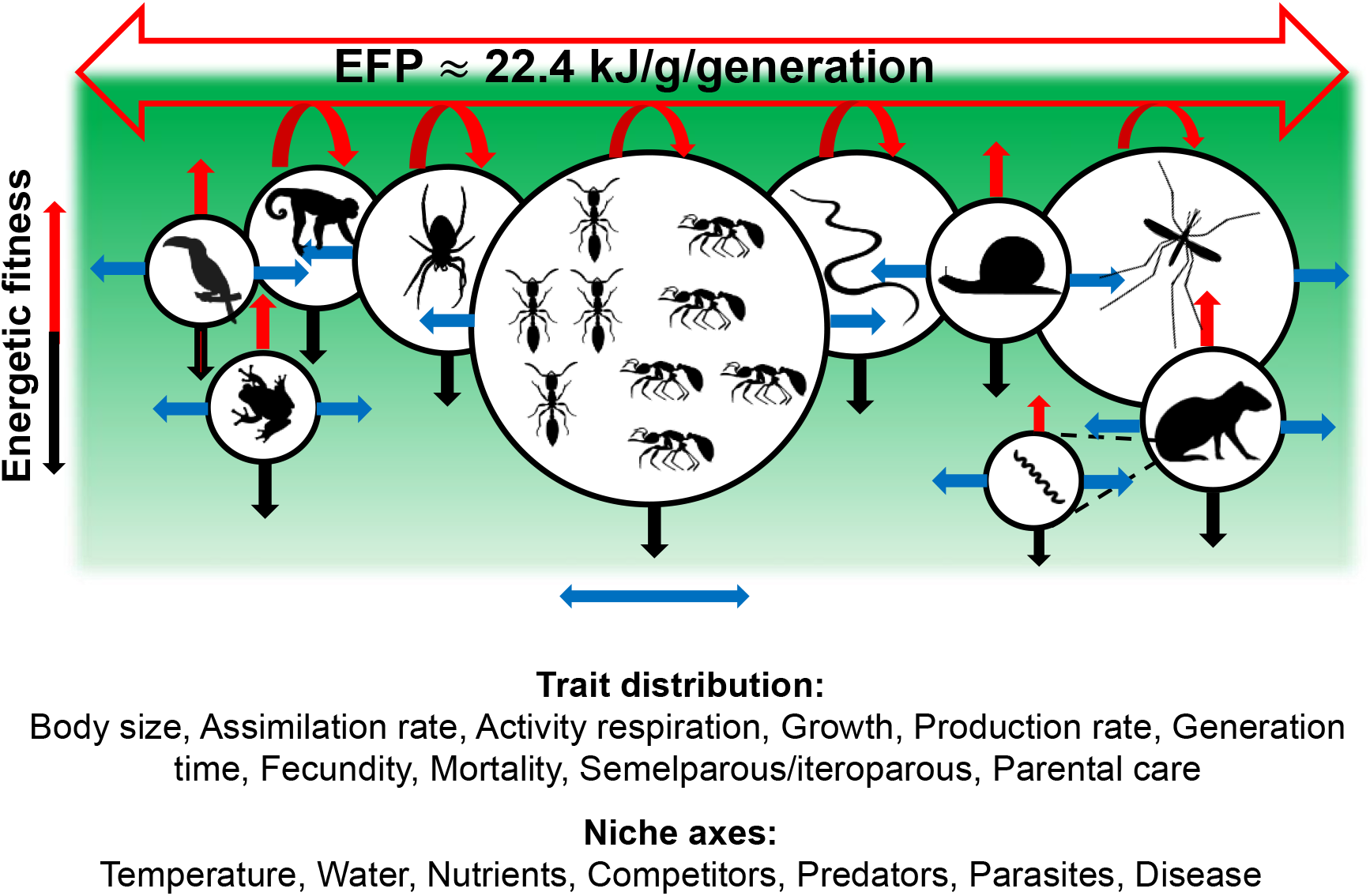
Schematic diagram representing diverse yet equally fit species. The joint effects EFP energetics and Red Queen interactions on the assembly of a hypothetical ecological community. As they compete for usable energy the coexisting species (represented by the circles) continually diversify in trait distributions and environmental requirements along niche axes (blue arrows). Some species are able to obtain a larger share of usable energy (size of circle), increase in abundance and energetic fitness (red arrows), and push up against the maximum steady state value of *E* = 22.4 kJ/g/generation. Other species are unable to keep up in the zero sum Red Queen game, decrease in abundance, and eventually go extinct (black arrows). Icons reused under Public Domain phylopic.org with special credits to Sarah Werning (Capuchin), Gareth Monger (Spirochete) and (Birgit Lang) Spider under https://creativecommons.org/licenses/by/3.0/

At this steady state the energetic fitness of each species is very close to the canonical 22.4 kJ/g/generation of the EFP (eq 1). This value is set by the biophysical laws of metabolism and life history, enforced by tradeoffs in physiology and behavior that limit the share of NPP that can be acquired by any given species and reinforced by the Red Queen zero sum interactions among species. As indicated above, the steady state hides complex non-equilibrium dynamics at smaller scales, where the component species fluctuate in abundance and their traits evolve adaptively in response to the abiotic environment and biotic interactions. When the environment changes and new resources become available, native species shift their niches or alien species colonize and exploit them. Some species are able keep up, but any increase in production and abundance is temporary, ultimately checked by ecological compensation and the Red Queen at the 22.4 kJ/g/generation steady state of the EFP. Other species are less successful; they fall behind in the zero-sum game, decrease in abundance and ultimately go extinct unless rescued by environmental change or new adaptations.

One consequence of these complex ecological and evolutionary dynamics is the enormous variation in the abundance, distribution and ecological niches of coexisting species. Darwin (1882, 6^th^ ed. p. 62) perceptively observed: “The most flourishing, or, as they may be called, the dominant species – those which range widely, are the most diffused in their own country, and are the most numerous in individuals….”. Since then many authors have addressed such variation, not only in abundance and distribution (e.g., Preston 1948; MacArthur 1957; Brown 1984, 1995; Williams 1964; Hubbell 2001; Harte 2011), but also in impacts on the physical structure, energy and nutrient flows, and species diversity of ecosystems (e.g., priority, pioneer, climax, foundational, engineer, keystone and apex predator species; e.g., Paine 1966; McNaughton and Wolf 1970; Dayton 1975; Lawton and Jones 1995; Schramski et al. 2015; Fukami 2015; Brandl et al. 2019; Enquist et al. 2020). This inequality in ecological relationships is another consequence of the EFP. Ecological communities are filled to capacity with nearly equally fit species, but their fitnesses are exactly equal only when populations are constant and natural selection is not operating. During departures from steady state, a small change in extrinsic environment or heritable trait can have a large effect on fitness and trigger large shifts in abundance, distribution, community composition and ecosystem processes. This interpretation is consistent with the well-documented “individualistic” responses of communities to climate change and invading species (e.g., Graham and Grimm 1990; Brown et al. 1997; Jackson and Overpeck 2000; Sax and Brown 2000; Williams et al. 2004; Sax et al. 2007).

Throughout its history, biology has focused on biodiversity – describing the unique anatomy, physiology, behavior and ecology of life stages and species, and organizing them into hierarchical groups based on shared features of structure, function and phylogenetic history. This endeavor has always included a healthy tension between empiricists, who focused on the variation and endeavored to quantify and organize it, and theoreticians, who focused on the repeated patterns and sought to “explain” them in terms of biophysical laws and mathematical equations.

This review and synthesis presents our vision for a still unfinished theoretical framework based on energy. We are indebted to pioneers who laid groundwork over a half-century ago: Boltzmann (1886), Thompson (1917), Lotka (1922), and Odum (1971). More recently, others been doing related work; some have contributed directly or indirectly to our work and others have challenged it. We cannot review or even cite them all here, but some common themes and still unresolved differences are in Box 5.

#### Box 5. Related work in metabolic ecology

##### 1) Scaling relations

As pointed out above, it has long been recognized that many traits scale predictably with body size and temperature. In the mid-1980s, following the pioneering studies of Kleiber (1932), Brody (1945), Hemmingsen (1960) and others, four important books attempted to synthesize the state of the science (Bonner and McMahon 1983; Peters 1983: Calder 1984; and Schmidt-Nielsen 1984, but see Heusner 1982, 1991). Each book presented a unique perspective, but there was consensus that metabolic rate and related processes scale as quarter-powers of body mass. But in the absence of a general theory, and interest ebbed. When West et al. (1997) purported to explain Kleiber’s law of *m*^3/4^ scaling of “metabolic rate’’ in terms of the structure and function of the fractal-like vascular networks that distribute metabolic resources, it was followed a flurry of supporting and critical studies. Points of contention included: i) vascular networks that violate critical assumptions of the WBE model (e.g., Bannavar et al. 2002, 2010; Chown et al 2007, White et al. 2011; Seymour et al. 2019; Aitkenhead et al. 2020); ii) variation in the parameters of fitted regression equations (e.g., Darveau et al 2002; Kozłowski et al. 2003; Kozlowski and Konarzewski 2004; Glazier 2005, 2010; Etienne et al. 2006; Apol et al. 2008; Dodds 2010); and iii) different statistical methods for analyzing data and evaluating hypotheses (e.g., Isaac and Carbone 2010; Kearney and White 2012; Uyeda et al. 2019; White et al. 2019). Many studies, and relevant theoretical and empirical issues, are addressed in Sibly et al. (2012).

In this paper, we circumvent many of these controversial issues by suggesting that the whole family of quarter-power scalings reflects the importance to fitness of generation time and the very general *m*^1/4^ scalings of biological times. We attribute this insight primarily to Linsdtedt and Calder (1981; see also Calder 1984). Like most ideas in science, however, it has its antecedents (e.g., Blum 1977; Bleuweiss et al. 1978; Richardson and Rosen 1979) and successors (e.g., Hainsworth 1981; Ginzburg and Damuth 2008; Colyvan and Ginzburg 2010).

##### 2) Rate of living

The EFP shows some resemblance to ‘rate of living’ or ‘pace of life’ theories (Rubner 1908; Pearl 1928). Recent versions posit that ageing is caused by energy metabolism: higher metabolic rates lead to shorter lifespans, because oxidative respiration generates free radicals and other byproducts that cause molecular and cellular damage and contribute to aging, senesce and mortality (for reviews and divergent assessments see Pearl 1928; Speakman 2005; Speakman and Król 2010; Selman et al. 2012; Hou and Amunugama 2015). It is becoming increasingly clear, however, that cause and effect are intertwined: metabolism does affect lifespan, but lifespan also affects metabolism, especially production. It takes longer to build and reproduce a larger, more complex organism, and so it must grow and resist aging and extrinsic mortality for longer than a smaller, simpler organism.

##### 3) Dynamic energy budgets

The body of work on dynamic energy budgets (DEB) by Kooijman and collaborators (e.g., Kooijman 1986; 2000; 2010; Nisbet et al. 2000, 2010, 2012; Sousa et al. 2008; 2010; Freitas et al. 2010; Maino et al. 2014) is a longstanding research program that paralleled work on a metabolic theory of ecology (MTE; Brown et al. 2004, 2018; Sibly et al. 2012a; Burger et al. 2019a). It may not be the “most comprehensive metabolic theory of life existing to date” (Jusup et al. 2017), but DEB has much in common with MTE. Both aim to elucidate fundamental rules of life based on mass-energy balance, other laws of physics and chemistry, and first principles and established facts of biology. Both provide an integrated framework of models and data. But there are differences in content and applications. DEB is more based on the biochemical and physiological details of metabolism, invokes some different assumptions, includes more parameters in its models, and has been more applied to practical problems of environmental policy and management. MTE has been more based on simple assumptions and models with fewer parameters, more focused on scaling relations and other phenomena of whole organisms, and more extended to address ecological and evolutionary patterns and processes at ecosystem, community, geographic and evolutionary scales. We do not, however, see any inherent conflict between DEB and MTE. The relative merits of alternative assumptions and models can be debated and resolved, and more and better data collected and compiled to test and extend the theories (e.g., Fig 3, 4 & Table 1; see also Marques et al. 2018). To a large extent, the perceived merits and demerits of the two theories are matters of subjective taste, not objective science.

### The Road Forward: Testing and extending the EFP

Recent decades have seen an explosion of interest in biological scaling and metabolic ecology. Regardless whether measured from the pioneering studies of Thompson, Lotka and Kleiber, the synthetic treatments of Peters, Schmidt-Nielsen, and Calder, or more recent work on metabolic scaling, DEB and MTE, the progress has been impressive. This broad research program has the common themes of theory based on first principles of physics, chemistry and biology, and empiricism based on compilation, analysis and interpretation of more and better data. On the one hand, there is optimism that these studies are leading to universal rules of life based linking the biological performance of individual organisms to their consequences for ecosystem ecology and global biodiversity. On the other hand, there is sobering realization of how much more work will be required to turn this vision into reality.

We do not presume to predict or prescribe the future. We do, however, provide our perspective on a few concrete steps that could be taken. Some of these are primarily theoretical. Progress will depend largely on extending the existing frameworks to develop models and make predictions about heretofore little explored phenomena. Others are more empirical. Progress will depend largely on applying new and better data to deductively distinguish between existing alternatives and inductively inspire unification and new ones.

#### A) Variations in life history

Both DEB and MTE rely on mass-energy balance to elucidate patterns and processes of life history. At present, however, there appears to be a gap between the “big picture” framework of MTE and EFP and the more detailed models (e.g., more parameters) of DEB. Neither theory has yet developed and tested models to address important aspects of such related phenomena as:

i. The “two-fold cost of sex”: Most animals reproduce sexually, so a female parent produces two offspring to replace herself and her mate in the next generation. Some of these animals have closely related parthenogenetic species: there is no sexual reproduction, offspring develop from unfertilized eggs, and a female parent produces only one female offspring than surviving to replace herself in the next generation. The resulting so-called “two-fold cost of sex” (e.g., Doncaster et al. 2000), expressed in terms of mass-energy balance, means that the lifetime cohort production, *P_coh_* in eqs 1 and 2 should be two times higher in a sexual species compared to an otherwise equivalent parthenogenetic one. Yet at steady state, both must have equal fitness. The apparent resolution to this puzzle is that it represents another example of ecological compensation. The two-fold advantage of a new parthenogen is only temporary; its population growth is ultimately checked by environmental limiting factors and other adaptive advantages of sexual reproduction and the resulting genetic variation come into play (see reviews by Doncaster et al. 2000; Lehtonen et al. 2012). The EFP and the pervasiveness of sexual reproduction implies that the energy cost and fitness benefit of sex is 22.4 kJ/g/generation, the “extra” energy required to produce a surviving male offspring.
ii. “Parental care”: DEB and MTE highlight insights gained by applying mass-energy balance and biophysical *dE/dt* currencies to analyses of life history. Most of the relevant fitness traits can be converted from the *dN/dt* variables of demography and population ecology to units of mass or energy. So, for example, it is relatively straightforward to define parental investment as the biomass or energy content of gametes plus any direct nutritional contribution prior to offspring independence (such as lactation in mammals and feeding in altricial birds). But there can be complications. An obvious example is the effect on energetic fitness of internal gestation and live birth in giving unborn offspring mortality similar to the mother (e.g., in sharks, mammals, and some reptiles, compared to teleost fish, amphibians, and birds). A more complicated problem is how to quantify the energetic fitness due to parents and members of social groups “teaching” and grooming offspring and protecting them from predators and parasites. Such parental care has seemingly been a major influence on allocation of production, values of *F* (1 – *F*), and evolution of life histories in taxa such as primates and mouth-brooding cichlid fish.
iii. Exceptions that prove the rule: Conceptual frameworks and formal models are often useful even when they fail: assumptions do not hold, additional factors or parameters are required, or data appear to refute predictions. As mentioned above, the steady state assumption of EFP is often violated, especially at small space and time scales. This does not invalidate the concept of energetic fitness; the mechanistic foundations of the EFP (eqs 1 and 2), and the prediction that ecological and evolutionary dynamics tend to result in nearly equal fitness across species (Fig. 2). It does, however, mean that models with more functions and parameters will likely be required in instances. It should be informative to test the EFP by applying it to exceptional cases, such as parthenogenetic species; parasites, insects and amphibians with complex life cycles; cyclical populations; and colonizing and declining species.
iv. Experimental tests: DEB, MTE and EFP share the key assumption of mass-energy balance. But like the Hardy-Weinburg equilibrium (HWE), their greatest contribution may be applications and extensions to predict dynamic responses when the steady state assumption does not hold. Examples from applied environmental sciences include harvested populations of game and ocean and freshwater fish, control of pests and colonizing invasive species, and management interventions to conserve endangered species and ecosystems. There are also opportunities to conduct controlled, manipulative experiments for direct tests of theoretical predictions using both wild animals and model laboratory organisms, such as rats and mice, fruit flies, the nematode *C. elegans*, and numerous unicellular eukaryotic and prokaryotic. There is a sizeable literature on experimental selection on *dN/dt* life history traits, but abundant scope for experiments to test effects of selection on the parameters of energetic fitness. The EFP predicts that selection to increase *P_coh_*, *G*, and *F* should result in increased energetic fitness. It is readily apparent that artificial selection on domesticated animals has resulted in substantial increases in rates of biomass production (growth and hence *P_coh_*) for human food consumption, and this has resulted in tradeoffs in traits that affect *G* and survival in the wild.

#### B) Biodiversity and applied ecology

The EFP has utility in understanding the general rules of life with applications to systems science, human and applied ecology based from first principles of physics, chemistry and biology and the compilation, analysis and interpretation of new data. These include:

i. *Body size spectra and energy equivalence*. – The empirical observation that in general small organisms are abundant and large ones are rare led to the concept of the biomass spectrum. Pioneering papers by Sheldon (Sheldon and Prakash 1972; Sheldon et al. 1977) showed that in pelagic marine ecosystems the number of individual organisms across all trophic levels scales approximately as the inverse of body mass, 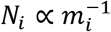, so the total biomass of the “particles” is independent of body size: 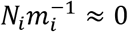 (e.g., Jennings 2005; Andersen et al. 2016). A related pattern within a trophic level is that population density scales as the inverse of Kleiber’s rule, so total rates of energy use (assimilation and respiration) are independent of body size: 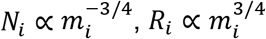, and 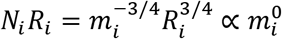 (e.g., Damuth 1981; White et al. 2007; Isaac et al. 2013).
ii. *Trait space*. – As shown in Fig. 4, the temporal sequence and quantitative allocations to ontogenetic growth and parental investment vary enormously across species, yet all of them have equal energetic fitness. It is obvious that these observed patterns are only a subset of the possible combinations, and most of them do not have equivalent fitness. This raises interesting questions about the trait space of life history parameters (e.g., Blonder et al. 2015; Morrow et al 2018) and how they interact with dynamic environments to form communities (e.g., Enquist et al. 2015; Burger et al. 2019b). How can they be combined, transformed, and reduced to a subset of traits arrayed along quantitative axes that accounts for most of the observed variation? Presumably such an analysis will reveal absolute constraints, relative tradeoffs, other important patterns and lead to insights into the biophysical underpinnings, ecological relations and evolutionary processes. It should help to answer questions such as: Why the strategy of producing enormous numbers of miniscule eggs that develop on their own and suffer enormous mortality has apparently facilitated the success of teleost fish as they invaded the oceans, diversified explosively and largely replaced sharks and rays after the K-T mass extinction? and How to explain the sometimes large differences in allocation to ontogenetic growth and parental investment by the two sexes of the same species (e.g., Sibly et al. 2012b).
iii. *Ecosystem energetics*. – MTE has been applied to link metabolism of individual organisms to the structure and dynamics of ecosystems, resulting in deductive tests of theoretical predictions and discovery of new empirical patterns. For example, body size and temperature play major roles in the structure of food webs (e.g., Brown et al. 2005; Schramski et al. 2015; Grady et al. 2019; Hatton et al. 2019). The EFP makes specific predictions linking energy flows from individual performance and population life history to ecosystem energetics. The parameter *F* and (1 – *F*) are closely related to the number and relative size of offspring. For example, mammals, which have a few relatively large offspring pass on more than half of cohort production to the next generation; whereas fish and invertebrates that produce large numbers of tiny offspring but lose more than 90% (sometimes much more than 99%) of cohort production to pre-reproductive mortality in the ecosystem (e.g., Brandl et al. 2019). These new insights from the EFP have implications for understanding ecosystem dynamics in space and time and the energetic ramifications throughout ecosystems during temporary departures from steady-state populations.
iv. *Human ecology*: DEB, MTE and their underpinnings in biophysical laws have been applied to human ecology – in particular, to correlates and consequences of energy use by humans over the historical transition from aboriginal hunter-gatherers in approximate steady state with their environment to the modern agricultural-industrial-technological societies whose unsustainable practices are transforming the biosphere (Burger et al. 2012). Interdisciplinary studies, at the interface between the biophysical science of ecology and the social sciences of economics and sociology, highlight the roles of energy and other resources on the growth, socioeconomic development, and environmental impacts of modern humans (e.g., Brown et al. 2011; Nikola et al. 2013; Syvitsk et al. 2020). A particularly interesting example is how the urban transition – the increasing concentration of humans in cities – can be understood in terms of a combination of biological constraints inherited from primate ancestors and modern technological innovations (e.g., Burger et al. 2017; Burger and Fristoe 2018; Burger et al. 2019c).

### Conclusion: Toward a modern ecological synthesis

The mid-20^th^ Century saw the development of the Modern Evolutionary Synthesis – a unified body of theory that incorporated the newly discovered biological laws of inheritance to explain patterns and processes of variation and change in living and fossil organisms. The Hardy-Weinberg Equilibrium, by assuming steady state and incorporating laws of Mendelian genetics, played a major role. Similarly, we see the Equal Fitness Paradigm, by assuming steady state and incorporating biophysical laws, playing a seminal role in a Modern Ecological Synthesis – a unified body of theory that explains patterns and processes of interactions between organisms and their environments in terms of the struggle for free energy. Such a theory has the potential to incorporate and integrate some of the most fundamental features of living things: i) metabolism, the intake, processing and expenditure of energy and materials by organisms; ii) scalings of physiological and life history rates and times with body mass and temperature; iii) universal features of demography and population dynamics; iv) flows of energy and materials through ecosystems; and v) origin and maintenance of biodiversity. Some of the seminal questions and predictions are given in Box 5. Much remains to be done to achieve unification, but the rate of recent progress provides grounds for optimism.

## Acknowledgements

This paper was outlined at Jim and Astrid Kodric-Brown’s home in Morro Bay over several days in January 2020, following presentations and discussions at the American Society of Naturalists meeting in Asilomar, California. JRB acknowledges funding from the Bridging Biodiversity and Conservation Science program at the University of Arizona and the National Science Foundation. We thank the Enquist lab group for discussions and Simon Brandl, Adam Chmurzynski, Matiss Castorena Salaks, and Richard Sibly for comments on the manuscript.

